# The “sex-specific effect:” Evaluating analytical approaches to sex-dependence in the behavioral and brain sciences

**DOI:** 10.64898/2026.02.04.703900

**Authors:** M. T. Olivier, A. W. Brown, S. Chung, C. J. Vorland, D. L. Maney

## Abstract

Detecting a sex difference in response to a treatment or intervention, often reported as a “sex-specific effect,” requires statistical comparison of the response across sex. Here, we investigated analytical approaches used to test for such effects in the behavioral and brain sciences. Of 200 recent articles containing terms such as ‘sex-specific’ or ‘gender-dependent’ in their titles, only 24% presented appropriate evidence supporting the claim: the effect was compared statistically across sex and results consistent with the claim were reported. In most articles (58%), no test was conducted that could have supported the title claim. Only 15% of studies on non-human animals supported the claim with appropriate evidence, which was significantly less frequently than studies on human participants (34%; *p* = 0.002). The use of appropriate analytical approaches was unrelated to journal rank or the citation impact of the article. We conclude that claims of sex/gender-dependent effects in the behavioral and brain sciences are only infrequently supported by appropriate evidence.

---

Advances in public health and medicine rest on rigorous research conducted on diverse populations. In recent decades, sex has been singled out as a critical dimension of this diversity, prompting policies and guidelines designed to increase the inclusion of females and males in biomedical research (CIHR, 2021; Heidari et al., 2016; 2024; NIH, 2015; Peters et al., 2021). An important rationale for such policies is that the routine exclusion of an entire sex or gender from research design, analysis, and reporting is difficult to reconcile with standards of scientific rigor or broad generalizability (Clayton & Collins, 2014; Health Canada, 2022; Heidari et al., 2016). Since the implementation of “Sex as a Biological Variable” (SABV) by the National Institutes of Health in the U.S. and “Sex and Gender Based Analysis” (SGBA) by the Canadian Institutes of Health Research (CIHR), the proportion of studies that include females and males has indeed increased (e.g., Kim et al., 2024; Mamlouk et al., 2020; Rechlin et al., 2022; Woitowich et al., 2020), suggesting that the policies are having a positive effect on sex inclusion.

Beyond concerns about generalizability, sex-inclusive research policies are grounded in the view that men and women differ systematically in patterns of disease, treatment response, and adverse drug effects, and that further differences are likely widespread but as yet undocumented (Clayton & Collins, 2014; Finn et al., 2025; Health Canada, 2022; Heidari et al., 2016). Since the SABV policy went into effect in 2016, reports of sex differences have increased sharply (Maney & Rich-Edwards, 2023). Although SABV and most similar policies do not require investigators to formally compare the sexes, the sexes are nonetheless compared more than 80% of the time and conclusions of difference are often highlighted in titles or abstracts (Finn et al., 2025; Garcia-Sifuentes & Maney, 2021).

## Box 1. What is the DISS error?

The “Difference in Sex-specific Significance” (DISS) error is an inappropriate analytical approach in which within-sex tests are conducted to determine whether males and females responded differently to a treatment (Maney & Rich-Edwards, 2023). In a design such as (a) below, the DISS approach does not compare the sexes directly; instead, *p* values are generated within-sex and, if they are discrepant with respect to statistical significance (b), the effect is declared to be sex-dependent. But the DISS approach does not test whether the effect depends on sex; the sexes have not been compared statistically. To test for a difference in the treatment effect, we could test for a sex by treatment interaction; a significant interaction would provide evidence that the effect differed between females and males.

The DISS approach dramatically reduces power to detect the effect of treatment, and biases researchers toward positive findings of “difference” (Bland & Altman, 2011). In the example below, the sexes appear to have responded similarly to the treatment, but the study was underpowered to detect the effect when the sample was limited to males only. DISS has thus led to a conclusion of difference when it is unlikely that the sexes responded differently. The DISS approach can also cause sex-dependent effects to be missed, for example when underpowered to detect an effect in the within-sex tests yet there was a significant sex by treatment interaction (Vorland et al., 2023). For more information about the error and sample data sets, see Rich-Edwards and Maney (2003).

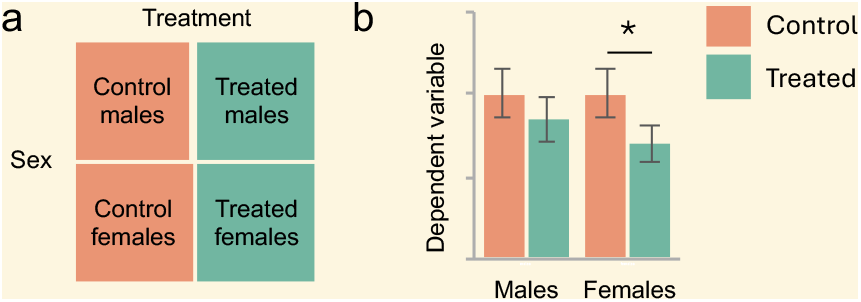

In the biomedical sciences, evidence for a sex difference requires a statistical comparison between the sexes. For sex-dependent effects, the comparison typically takes the form of an interaction term—it explicitly tests whether the effect of interest depends on sex (Chin & Christians, 2015; Eliot et al., 2023; Gelman & Stern, 2006; George et al., 2016; Karp, 2025; Makin & de Xivry, 2019; Nieuwenhuis et al., 2011; Phillips et al., 2023; Rechlin et al., 2022; Rich-Edwards & Maney, 2023; Sainani, 2010; Vorland, 2021). Training materials provided by organizations such as CIHR and NIH, although they emphasize generalizability, provide incomplete, misleading, or incorrect instruction on analysis of sex-based data (Gompers et al., 2024; see also Chiocca & Maugeri, 2020; Heidari et al., 2016). For example, these trainings each endorse what has been called the ‘difference in sex-specific significance’ (DISS) error (Maney & Rich-Edwards, 2023), which is the practice of testing effects separately within females and males and then inferring a “sex-specific effect” when the effect is statistically significant in one sex but not another (Box 1). Approaches that do not compare the effect statistically across sex do not test for a sex-dependent effect.

## Box 2. Terminology

**Sex and gender:** The terms ‘sex’ and ‘gender’ emerged from different scholarly traditions and are not equivalent (Maney et al., 2025). Their operationalization can vary according to research question and other contextual factors (Pape et al., 2024). Here, we use the terms to refer to binary categories to which animals or participants were assigned in the articles in our sample. We investigated how these categories are incorporated as variables into statistical models; because no study in our sample distinguished between sex and gender, treating them as equivalent in this case does not result in loss of generality. We use the term ‘sex’ to refer to sex or gender, ‘males’ to refer to males or men, and ‘females’ to refer to females or women. None of the articles in our sample clearly operationalized any of these terms and in all cases the categories were binary.

### Effect

We adopt the term ‘effect’ to refer to the influence or association of a non-sex factor that has been tested in males and females. ‘Effect’ is commonly used in this context, regardless of whether an investigation is causal or associative in nature. For example, a ‘sex-dependent effect’ could refer to the effect of an experimental factor that depends on whether mice are female or male; it could also refer to an association between two variables, such as a risk factor and a health outcome, that differs between men and women.

### Behavioral and brain sciences

Our goal was to sample broadly from literature relevant to behavior and the brain. Our corpus consisted of articles tagged in Web of Science with the following research areas: behavioral sciences, clinical neurology, neurosciences, psychiatry, psychology, and substance abuse. We refer to these research areas as our ‘focal’ areas. See Methods for a complete list of all related research area tags.

Sex differences relating to the brain, cognition, and behavior have attracted attention for centuries (Maney, 2014; 2016). Such differences can be highly contentious (Cahill, 2014; DeCasien et al., 2022; Eliot, 2024; 2025) and are often misrepresented (Casto & Maney, 2024; Fine, 2012; Fine et al., 2019; Maney, 2014; 2016; Rippon et al., 2021; Sanchis-Segura et al., 2026; Vorland et al., 2023). Because sex differences are frequently emphasized and culturally salient in the neurosciences specifically (Fine, 2012; Jordan-Young & Rumiati, 2012), it is especially critical for neuroscientists to maintain high standards of scientific rigor when testing whether females and males respond differently to an exposure or intervention. Here, we examined the analytical approaches used to test for sex-dependent effects in the behavioral and brain sciences (see Box 2). We analyzed 200 recently published articles featuring claims of a sex-dependent effect in the title, indicating that it was a main finding. These articles therefore represent a corpus in which we should expect explicit and convincing statistical evidence supporting the claim.

## Results

### Article characteristics

To identify articles focused on sex-dependent effects, we searched for claims that males and females (see Box 2) responded differently to a treatment, exposure, intervention, or other non-sex factor. We use the term ‘treatment’ here to refer to this non-sex factor, with the caveat that it could have been a variable such as genotype, age, season, presence of a health condition, etc. We searched the Web of Science Core Collection (Clarivate Analytics) for articles with terms such as ‘sex-specific’ or ‘gender-dependent’ in the title (Fig. 1a), restricting the results to articles published between 2019 and 2023 in fields related to the brain and behavior (Box 2). Using a weighted sampling algorithm (Chung et al., 2025), we selected 200 articles from the resulting 1,341 (see Methods, Table S1, Supplementary Data File 1). The most common search term to appear in the titles was ‘sex-specific,’ followed by ‘sex-dependent’ and ‘gender-specific’ (see Fig. 1a). Half of the articles (n = 100) involved human participants (Fig. 1b); of these, four also involved mice. Among the 100 articles on only non-human models, most (n = 95) were on rodents.

**Figure 1.**
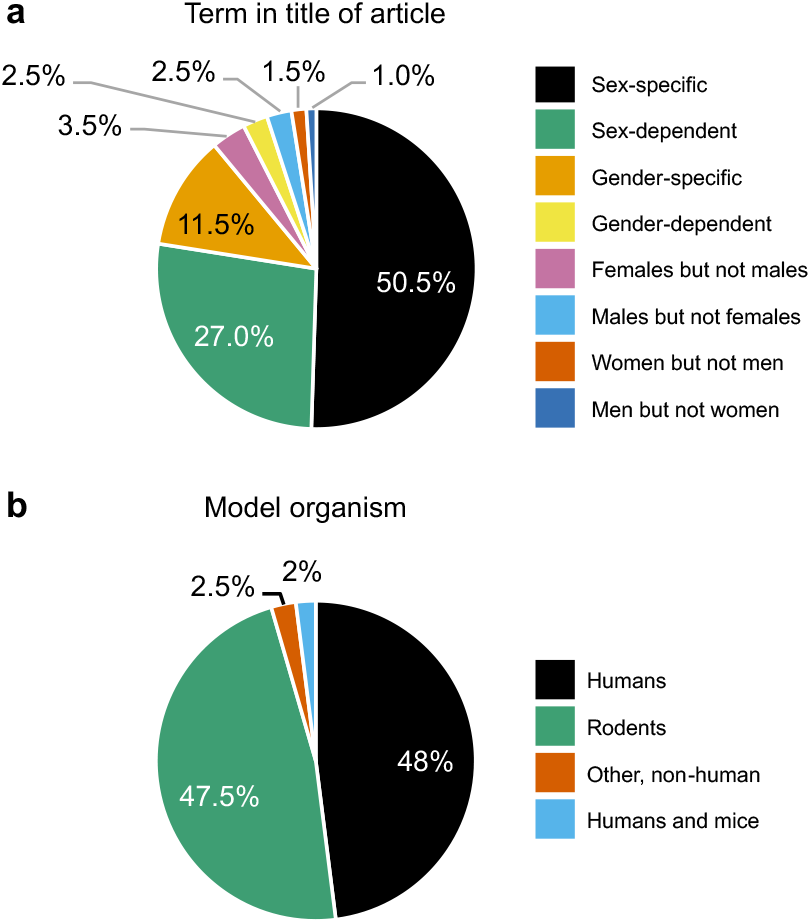
The use of sex-specific terms and model organisms in a sample of 200 articles in the behavioral and brain sciences. (a) shows the percentages of articles with each of our search terms in the title. (b) shows the model organism in each of the articles. Non-human rodent models included mice (n=51), rats (n=35), both rats and mice (n=1), prairie voles (n=3), Siberian hamsters (n=2), degus (n=1), gerbils (n=1) and red squirrels (n=1). The remaining five articles were on birds (collared flycatchers, n=1; blue tits, n=1), Barbary macaques (n=1), pigs (n=1), and *Drosophila* (n=1).

We treated ‘sex’ and ‘gender’ as equivalent for the purposes of article inclusion and coding (see Garcia-Sifuentes & Maney, 2021) and use the term ‘sex’ throughout (Box 2). None of the articles independently incorporated both sex and gender into statistical analyses.

### Analytical approaches

Our main goal was to document the types of evidence used to support claims of sex-dependent effects. We classified analytical approaches using criteria described in the Methods and Table 1; distributions are shown in Figure 2. Of the 200 articles, 48 (24%) supported the sex-related title claim with appropriate evidence, meaning that the effect mentioned in the title was statistically compared across sex and the result of that comparison supported the claim (see Table 1). In most of these (n = 39) a multifactorial ANOVA was performed with sex as a factor and a statistically significant interaction was reported between sex and the other factor(s). Seven articles reported a statistically significant difference between within-sex associations (e.g., within-sex Pearson correlations were compared with each other), and two compared other effect estimates, such as odds ratios, across sex.

**Table 1.** Codes used for categorizing articles according to analytical approach.

| General analytical approach | Presentation of evidence | Evidence designation for graphs |
| --- | --- | --- |
| **1. Claim in title was tested statistically (e.g., in a multifactorial ANOVA with sex as a factor, comparison of associations, etc.)** | 1.1. Authors reported a $p$ value indicating a significant interaction (a), significantly different within-sex correlation (b), or other appropriate statistical evidence (c) supporting the claim in the title | Appropriate evidence |
| | 1.2. Authors reported a significant interaction with sex but did not report the relevant statistic (e.g., $p$ value) | Missing evidence |
|  | 1.3. Authors claimed to have performed an appropriate test but did not report the result or state statistical significance, either significant or non-significant |  |
| | 1.4. Authors reported a $p$ value indicating a non-significant interaction | Contradicting evidence |
| | 1.5. Authors reported that the interaction was not significant but did not report the $p$ value | |
| **2. Claim in title was not tested statistically** | 2.1. Authors performed within-sex tests only (DISS error) | No direct test of claim in title |
|  | 2.2. Authors tested whether sex explained variation in the overall model (e.g., main effect), then performed within-sex analyses |  |
|  | 2.3. Authors performed no quantitative comparisons |  |

*Note: During the coding process, we also coded whether authors 1) conducted post-hoc tests regardless of whether the interaction was significant or not significant, and/or 2) conducted sex comparisons within treated groups, the results of which are described in the text narrative.*

**Figure 2.**
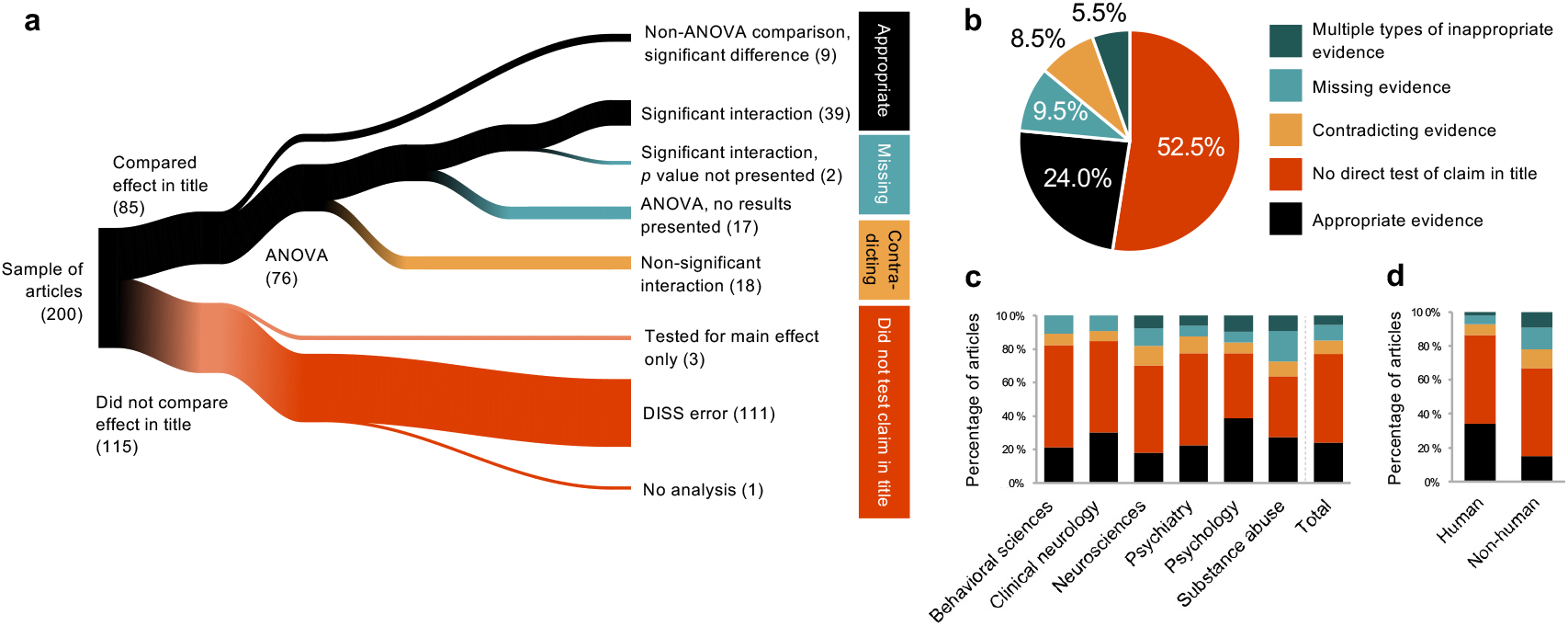
Analytical approaches supporting claims of sex-dependent effects in the titles of articles in the behavioral and brain sciences. The river plot (a) depicts the numbers of articles with each type of evidence. The approaches and colors shown on the right side of the panel correspond to those in the other panels (see Table 1 for categories and descriptions). In (b), articles in which authors used multiple types of inappropriate evidence (n=11) are shown in dark teal (DISS and missing ANOVA results, n=6; DISS and contradicting ANOVA results, n=4; missing ANOVA results and contradicting ANOVA results, n=1). The 11 articles with multiple types of inappropriate evidence are included in the river plot (a) as DISS (n=10) or contradicting (n=1). (c) Across the six focal research areas, articles in the field of psychology were the most likely to contain appropriate evidence of the claim of sex-dependent effects in the title, and those in neuroscience the least likely, compared separately with all other areas. Articles that were tagged with two research areas, for example neurosciences and behavioral sciences, are included in both relevant columns. (d) Studies with human participants were significantly more likely to include appropriate evidence than those with non-human animals (*p* = 0.002).

In the remaining 152 articles (76%), evidence used to support the title claim did not meet criteria for appropriateness in that a statistical sex comparison was claimed to have been conducted but the results were missing, or the sex comparison was missing entirely. Thirty-seven of these took a multifactorial ANOVA approach; of these, two claimed a statistically significant interaction between sex and the other factor(s) but no *p* value was reported for that interaction. Seventeen articles claimed to have performed a multifactorial ANOVA with sex as a factor, but failed to report both the *p* value and the result (significant or not) of the interaction test. Together, these 19 articles missing the *p* value, the result, or both were coded as ‘missing’ (Fig. 2). In 18 articles, the interaction was explicitly reported as not statistically significant, indicating that the claim in the title was not supported by the data and that the null hypothesis — that the effect in question did not depend on sex — should not have been rejected. For these articles, 15 of which included the non-significant *p* values for the interaction, the ANOVA results therefore contradicted the claim in the title. These articles were coded as ‘contradicting’ (Fig. 2).

In the 76 articles using ANOVA, whether classified as appropriate or inappropriate, we noted whether post-hoc tests of the effect were conducted within sex. Post-hoc tests are typically predicated on first detecting a significant interaction, which provides evidence that the effect differs by sex (see Rich-Edwards & Maney, 2023). In 22 of the 76 articles, it appeared that all ANOVAs were followed by post-hoc within-sex tests regardless of whether the interaction with sex was statistically significant, suggesting that the interaction test from the ANOVA had been disregarded. Most of these articles were ultimately coded as ‘inappropriate,’ but two were coded as ‘appropriate,’ because the title claim happened to align with significant interactions.

In most articles (58%), no test capable of supporting the title claim was conducted. These were coded as ‘no direct test.’ In 111 of these 115 articles (56% of the 200 articles in the sample), the effect in the title was tested separately in females and males, and discrepant *p* values were interpreted as a sex difference (the DISS error, see Maney & Rich-Edwards, 2023; Box 1). In two of these articles, sexes were compared within treated groups, disregarding the control groups. This error, which is similar to DISS, has also been reported to occur in sex differences research (Garcia-Sifuentes & Maney, 2021; Phillips et al., 2023).

In three of the 115 articles in which the effect was not compared across sex, sex was included in an omnibus test prior to within-sex analysis, but that model did not test for sex-dependence of the effect in the title. Instead, the model tested for a sex difference in the outcome measure regardless of the other factors (e.g., main effect of sex), meaning that a significant sex difference did not relate to the claim in the title. The last of the 115 articles contained no quantitative analyses (see Figure 1a).

### Research area

We next investigated the analytical approaches used within each of our focal research areas (Fig. 2c) as defined by the Web of Science tags (see Methods). In our sample, psychology had the highest percentage of articles with appropriate statistical evidence (12 out of 31 articles, or 38.7%). The neurosciences had the lowest percentage of articles with appropriate statistical evidence at only 18.2% (14 out of 77 articles). In an exploratory analysis, we investigated the probabilities of using appropriate evidence in each research area individually, compared with all other areas; for example, we compared the set of articles tagged with neuroscience with those not tagged with neuroscience, and so on for each research area. In a model that included all six research areas, the set tagged with psychology was statistically significantly different from the non-psychology articles (OR = 4.705, 95% CI: [1.255, 18.127], *p* = 0.022), indicating that psychology articles were significantly more likely to contain appropriate evidence for the claim in the title than articles outside psychology. We found no other significant differences when comparing each area against all other areas (see Tables S2 and S3 for complete results). We did not conduct pairwise comparisons between the areas because some articles were tagged with two research areas. We therefore do not draw conclusions about whether any research area was more or less likely to contain appropriate evidence than another specific area.

Because we were particularly interested in tracking DISS errors, we examined their prevalence across the research areas. Overall, the DISS approach was common in all six areas (Fig. S1a). In our sample, psychiatry articles were most likely to feature a DISS approach to the title claim, at 61.2% (30 out of 49 articles) and substance abuse the least, at 40.9% (9 out of 22 articles). As above, we compared the prevalence of the error between the set of articles with a particular research area tag and those without it. These tests showed differences that were statistically significant for psychology (OR = 0.271, 95% CI: [0.082, 0.869], *p* = 0.030) and substance abuse (OR = 0.227, 95% CI: [0.060, 0.824], *p* = 0.026) (Fig. S1a), indicating that articles in each of these areas were less likely to employ a DISS approach to support the title claim than articles outside these areas (see Tables S4 and S5 for complete results). Again, we did not conduct pairwise comparisons between research areas, so we do not claim that the DISS error is significantly more or less prevalent in any one area than in another.

### Human vs. non-human models

Next, we investigated whether the appropriateness of analytical approaches was related to the choice of model organism (Fig. 2d). To perform this analysis, we excluded the four articles on both humans and mice. Of the 96 articles that included only human participants, 33 (34.3%) provided appropriate statistical evidence to support the claim in the title, whereas only 15 of 100 articles (15%) on non-human animals did so (non-human vs. human OR = 0.337, 95% CI: [0.165, 0.663], *p* = 0.002; Figure 2d; see Tables S6 and S7 for complete results).

The DISS error was the most common analytical approach, both within the set of articles on human participants and in those on non-human animals. Of the 96 articles that included only human participants, 49 (51%) took a DISS approach to the title claim and 59 of the 100 articles (59%) on non-human animals did so. The odds of taking a DISS approach were 1.380 times higher for articles on non-human animals than for articles on human participants (95% CI: [0.786, 2.435]), but this difference was not statistically significant (*p* = 0.263) (Figure S1b; see Tables S8 and S9 for complete results).

### Prestige and impact

We tested whether the choice of an appropriate analytical approach was associated with the prestige of the journal or the impact of the article itself. We operationalized prestige as the journal’s quartile from Journal Citation Reports, which is based on the impact factor of the journal (see Methods). We found no association between the appropriateness of an article’s analytical approach and the quartile of the journal that published it (Fig. 3a; Reference group: Appropriate; OR = 1.028, 95% CI: [0.486, 2.177], *p* = 0.942). We also found no association between the use of the DISS approach and the quartile (Reference group: no DISS; OR = 0.836, 95% CI: [0.435, 1.607], *p* = 0.590; Fig. S2a; see Tables S10 and S11 for complete results). These findings indicate no compelling evidence that prestigious journals are significantly more or less discerning when it comes to rigor in the analysis of sex-based data than their less prestigious counterparts.

**Figure 3.**
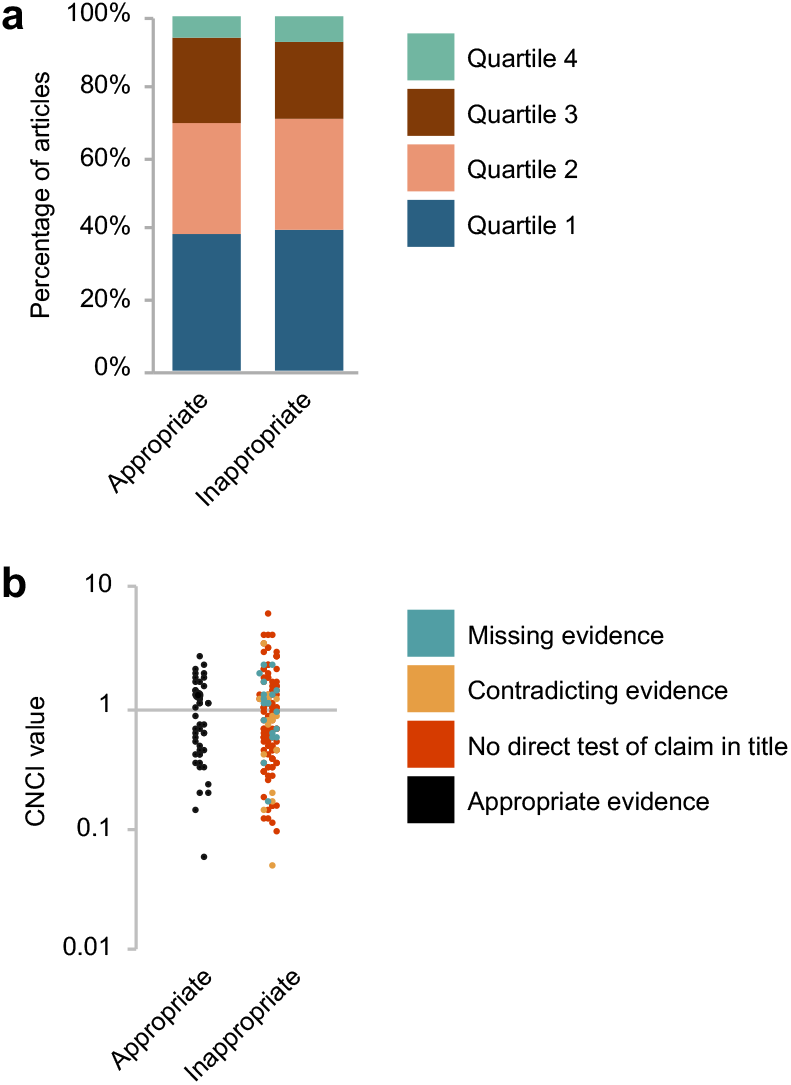
Associations between journal quartile or article impact and choice of analytic approach. The appropriateness of the analytic approach was associated with neither (a) journal quartile ranking from Journal Citation Reports (*p* = 0.942) nor (b) Category Normalized Citation Impact (CNCI) score from InCites (*p* = 0.859). Quartile ranking indicates the relative Journal Impact Factor within a research area, with Q1 being the highest. CNCI values greater than 1.0 indicate above-average citation impact, whereas values less than 1.0 indicate below-average citation impact.

We operationalized the impact of the article using the Category Normalized Citation Impact (CNCI) score, an article-level citation metric provided by InCites (Clarivate Analytics). This metric represents the ratio of an article’s observed citation count to the expected citation count for publications of the same document type, publication year, and Web of Science subject category (Clarivate, 2025). We found neither an association between the appropriateness of the analytical approach and the article’s CNCI (Fig. 3b, 2.6% increase in ‘inappropriate’ compared with ‘appropriate,’ 95% CI: [-25.8, 30.9]%, *p* = 0.859), nor an association between the use of the DISS approach specifically and the article’s CNCI (Fig. S2b; 2.9 % increase in ‘DISS’ compared with ‘no DISS,’ 95% CI: [-27.3, 21.5]%, *p* = 0.816; see Tables S12 and S13 for complete results). These findings suggest that articles with appropriate evidence for claims of sex-dependence are not significantly more or less impactful than articles without appropriate evidence.

### Claims of sex-dependent effects over the past two decades

Our findings of low rates of appropriate evidence for sex-dependent effects led us to ask about the extent to which such effects are being emphasized in different research areas. We plotted the number of articles in our six focal areas (see Box 2) that contained, in their titles, one of our search terms (e.g., “sex-specific”) spanning 2005 – 2025 (Fig. 4). For comparison, we did the same for 55 other research areas not focused on the brain, cognition, or behavior (see Table S3 for the complete list of research areas). The annual number of articles with title claims of sex-dependent effects in neuroscience, normalized to the total number of articles published in all 61 research areas each year, was generally higher than in the other areas and increased by more than five-fold over the 20-year period. In 2025, there were 2.5 times as many articles with claims of sex-dependence in the title in neuroscience than in the next-highest area, psychiatry.

**Figure 4.**
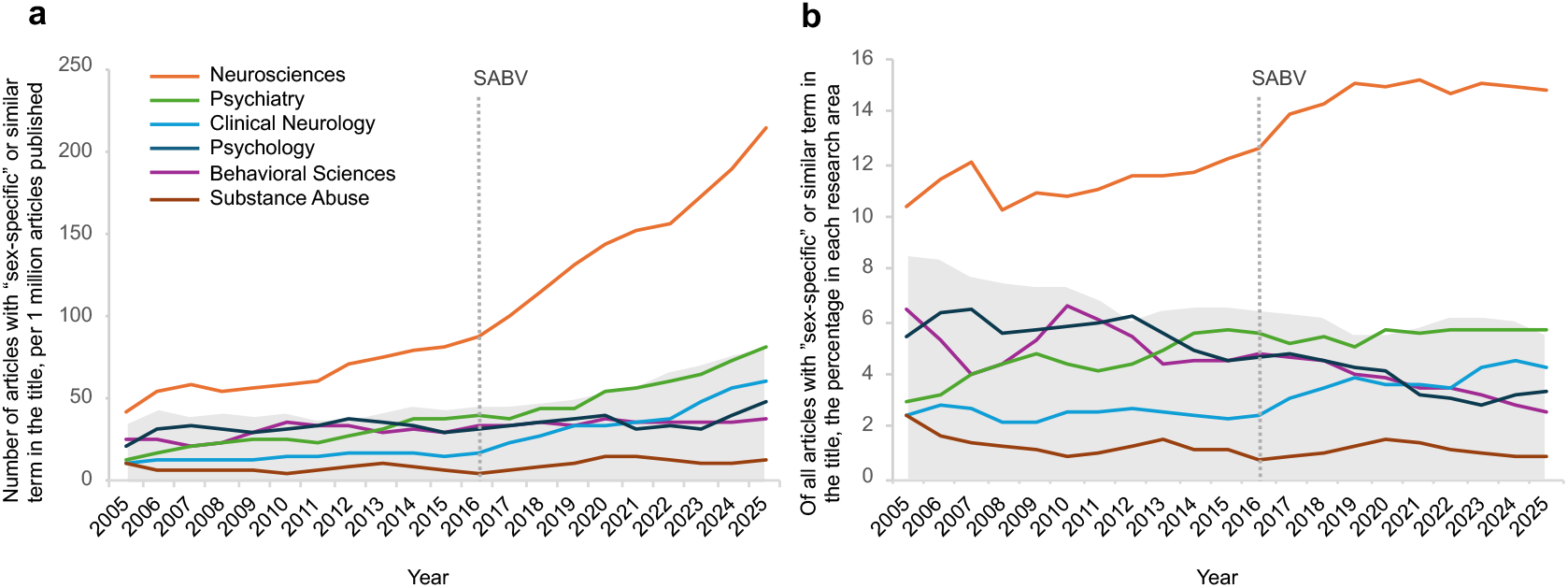
Title claims of sex-dependence over time. Each line represents articles with “sex-specific” or a similar term (see Fig. 1a) in the title for each of our six focal research areas. The gray area in the background is the range each year for 55 other research areas in Web of Science that do not focus specifically on the brain or behavior (e.g., radiology, allergy, surgery; see Table S3 for a complete list). (a) shows the number of articles each year, normalized to the total number of articles published in all 61 research areas to control for a general increase in publication rates over time. (b) each line shows, for any given article with “sex-specific” or similar term in the title, the probability that the article was tagged with that research area. For both graphs, we applied a 3-year centered moving average to reduce visual noise. All searches were conducted in January 2026, so the data for 2025 may underestimate publication counts. Source data are in Supplementary Data File 2.

To further visualize interest in sex-dependent effects in neuroscience compared with the other areas, we also plotted the probabilities that an article with a claim of sex-dependence in the title would be tagged with each of the 66 research areas (Fig. 4b). Consistently across the entire 20-year period, an article with such title claims was more likely to be a neuroscience article than any other area. By the time SABV was implemented in 2016, an article with a claim of a sex-dependent effect in the title was already twice as likely to be a neuroscience article than an article from the next most common area. By 2025, the chance had increased to three times as likely.

## Discussion

We report here that when articles in the behavioral and brain sciences include claims of sex-dependent effects in their titles, fewer than one-quarter (24%) present appropriate statistical support for that claim. We focused intentionally on claims made in titles because we expected those claims to be supported by rigorous statistical tests and convincing evidence. Yet, the overwhelming majority of these articles did not contain appropriate support for the claim. Our data reveal a failing of scientific rigor in the face of mandates that were implemented explicitly to improve it.

Claims of a sex-dependent effect must be supported with a statistical test of whether the effect depended on sex. In many articles in our sample, an appropriate test was performed but the result was not regarded as relevant to the claim; authors proceeded to post-hoc within-sex tests apparently without recognizing that the interaction test, not the post-hoc tests, is the test for sex-dependence. One article stated, “Post-hoc Tukey’s tests were done for all analyses to illustrate the interaction between groups.”

In the majority of articles in our sample (56%), males and females were not compared statistically at all. Instead, independent tests were performed within each sex, and discrepant *p* values were interpreted as a sex difference despite the absence of a formal comparison across sex (DISS error, Box 1). Almost all articles containing no sex comparison nonetheless included statements^1^ such as “The treatment induced deficits selectively in male offspring” (the article contained no test of whether males and females responded differently to the treatment); “Our results illustrate the importance of examining interactions between sex, behaviors, and genes” (no such interactions were formally tested); “To our knowledge, we are the first to describe sex-specific effects” (no test for sex-specific effects was conducted).

Two recent, smaller studies (Garcia-Sifuentes & Maney, 2021; Maney et al., 2023) also found high rates of inappropriate approaches to sex-based data. Because those studies were not focused exclusively on the behavioral and brain sciences, it was not possible to determine the prevalence of problems such as DISS errors in those research areas specifically. Here, we show that inappropriate approaches are widespread in research related to the brain and behavior, including in the neurosciences. Of the six research areas we included, articles in the neurosciences were least likely (18%; Fig. 2c) to test the claims of sex-dependence using appropriate approaches. This finding is particularly important in light of the abundance of articles in the neurosciences that contain such claims, compared with other fields. The number of neuroscience articles claiming sex-dependent findings in the title has been higher than in any other research area since at least 2005, and is currently two and a half times higher than in the next highest research area, psychiatry (Fig. 4). Our analysis of analytical approaches suggests that, on average, fewer than one in five of these claims in neuroscience is supported by appropriate quantitative analyses. Because DISS errors in particular cause strong bias toward false positive “findings” of difference (Bland and Altman, 2011; Brookes et al., 2004; George et al., 2016), the high rate of inappropriate approaches in neuroscience (82%) could be at least partially responsible for the high number of positive findings.

The prevalence of appropriate approaches in neuroscience (18%) was less than half that in psychology (39%). In psychology, null hypothesis significance testing (NHST) has long been treated as a framework for statistical inference, with explicit attention to matching statistical tests to the claims being made and to the interpretation of comparisons among effects (e.g., Cohen, 2008; Cohen, 1994; Cumming, 2013). In neuroscience, by contrast, several analyses have documented widespread inferential errors under NHST that arise not from the absence of statistical testing, but from failures to test precisely the claims being made (Nieuwenhuis et al., 2011), faulty interpretations of underpowered studies (Button et al., 2013), and over-reliance on dichotomous interpretations of *p* values (Hupé, 2015). These critiques suggest that in neuroscience, NHST is most often used to validate the presence or absence of effects rather than to test inferential claims about differences between effects. But NHST cannot logically be used to test for the absence of an effect, a practice that leads to invalid comparison between its presence and absence (Chin & Christians, 2015; Gelman & Stern, 2006; Maney, 2016; Makin & de Xivry, 2019; Sainani, 2010). It is this invalid comparison that directly underpins the DISS error, a specific case of the ‘differences in nominal significance error’ (Bland & Altman, 2011; George et al., 2016; Maney & Rich-Edwards, 2023; Rich-Edwards & Maney, 2023; Vorland, 2021).

We noted low adoption of appropriate approaches specifically in non-human animal research (15%). The higher rate of appropriate approaches in human research (34%) could be related, again, to the conceptualization of NHST in fields such as psychology, which is more likely to focus on human participants. Notably, training materials accompanying the SABV mandate, which targeted “preclinical” (non-human animal) research, endorsed the DISS approach (Gompers et al., 2024). Studies on non-human models typically have lower power than human studies (Button et al., 2013), increasing the risk that within-group analyses will fail to detect a true effect in one of the groups (Bland and Altman, 2011; Brookes et al., 2004; George et al., 2016), leading to a false perception of a sex-dependent effect.

We found no evidence that a lack of appropriate evidence for title claims prevented articles from being accepted to prestigious journals, as defined by impact factor quartile, or otherwise affected the articles’ impact, as assessed by a citation metric. Articles with inappropriate approaches were just as likely to appear in top-ranked journals (Fig. 3a) or to be cited (Fig. 3b) as those with appropriate evidence. Journals are currently under pressure to comply with the SAGER guidelines, which recommend that all data be analyzed and presented disaggregated by sex (Heidari et al., 2016; Van Epps et al., 2022). Thus, as is the case for the SABV policy, there is tension between compliance with the guidelines and maintaining rigor with respect to hypothesis testing, analytical approaches, and data interpretation. This tension, together with the ever-increasing popularity of sex differences in the brain (Fig. 4; see also Maney, 2016; Maney & Rich-Edwards, 2023), could at least partly explain why even prestigious journals may not require rigorous testing of “sex-specific” effects.

The consequences of disaggregating data and conducting separate tests go beyond invalid sex comparisons. First, the practice results in a profound loss of power due to halving the sample size of each test. Yet, some authors of articles in our study believed the DISS approach *retains* power; one article stated that “Data were analyzed separately for each sex to retain adequate statistical power to identify the within-sex effects.” Power to detect the main effect of interest is higher when all participants/samples are included in a *single* model that includes sex; this single model can also test whether the effect depended on sex (Karp, 2025; Nieuwenhuis et al., 2011; Phillips et al., 2023; Rich-Edwards & Maney, 2023). Second, the act of testing females and males separately itself increases the perception of difference (Saguy et al., 2021) as well as the chances of finding a spurious one. Within-sex tests can shift the focus of a project to differences that were not hypothesized in the first place. In at least two articles in our sample, the authors stated explicitly that they had no *a priori* hypotheses about sex. In others, the sex difference was rather buried, emerging more as an incidental finding than a major one. Yet, in each case, the finding seemingly shifted from exploratory to title-worthy (see Rich-Edwards & Maney, 2023).

The most serious consequence of claiming sex differences without evidence is the risk to equitable access to healthcare. In our sample of articles, calls for sex-specific diagnostic practices and medical treatments were widespread. For example, authors called for the development of different suicide prevention programs for men and women, different thresholds for substance use intervention, different cutoff scores when assessing risk for childhood conduct disorder, and different treatments for stress-related psychiatric disorders like anxiety and depression. None of these calls were supported by statistical sex comparisons. The likelihood that any effect in biomedicine is truly “specific” to only one sex is vanishingly small (e.g., Finn et al., 2025; Wallach et al., 2016). Recommending different treatments for men and women, without showing large, convincing sex differences in the effects of those treatments, could lead to needless and discriminatory barriers for patients of either sex, preventing them from receiving the most beneficial interventions (Pape et al., 2024; Rushovich, et al., 2023; Zhao et al., 2023).

In all the articles in our sample, sex was treated as a binary categorical variable, which, together with the conventions of NHST, constrains the questions we can ask, how we interpret our results, and the communication of findings. Opportunities to study sex-related variation in ways that move beyond simple categorical comparisons are increasing, however (see McLaughlin et al., 2023; Sanchis-Segura et al., 2026; Sanchis-Segura & Wilcox, 2024; Yang et al., 2025). Alternative strategies include modeling sex-related traits as continuous variables with bimodal distributions, representing sex as a multivariate constellation of characteristics, or adopting more exploratory frameworks such as discriminant analyses or probabilistic models that combine multiple sex-related measures into higher-dimensional phenotypic representations. By reducing reliance on *a priori* division of samples into two discrete groups, these approaches can support more nuanced understandings of biological variation while mitigating the tendency to over-binarize not only sex category, but also the statistical significance of effects within-sex: “yes” or “no.”

The primary stated goal of sex-inclusive research policies and guidelines is to improve not only generalizability but also rigor and reproducibility (Clayton & Collins, 2014; Health Canada, 2022; Heidari et al., 2016). Although more studies are now including females and males, our data show room for improvement with respect to the analysis and reporting of sex-disaggregated data. Efforts to uncover meaningful sex differences, absent analyses that produce meaningful evidence, risk producing a “literature of contradiction” (Rich-Edwards et al., 2018) muddling further progress and reducing the value of the policies. Sex-inclusive neuroscience will fulfill its promise only when inclusion is matched by analytical rigor.

## Methods

### Article selection

On May 24, 2024, we searched the Web of Science Core Collection for publications with titles containing phrases explicitly asserting sex- or gender-contingency. Our search string was: TI=(“sex-specific” OR “sex-dependent” OR “gender-specific” OR “gender-dependent” OR “males but not females” OR “females but not males” OR “male but not female” OR “female but not male” OR “men but not women” OR “women but not men”). These search terms were selected *a priori* to maximize specificity rather than exhaustive coverage. We restricted results to articles published between 2019 and 2023 and excluded meeting abstracts, proceedings papers, review articles, early access papers, book chapters, editorial materials, and news items. This process yielded 4,870 articles.

Next, we filtered the results to include only articles in the behavioral and brain sciences. In Web of Science, each academic journal is tagged with at least one research area, which is assigned to all the articles published in that journal. We selected the following Web of Science research areas to include in our analysis: behavioral sciences, clinical neurology, neuroimaging, neurosciences, psychiatry, psychology, applied psychology, biological psychology, clinical psychology, developmental psychology, educational psychology, experimental psychology, mathematical psychology, multidisciplinary psychology, psychoanalysis psychology, social psychology, and substance abuse. Several major journals that publish articles across disciplines, such as Nature, are tagged as “multidisciplinary sciences” or “medicine, general internal” on Web of Science. To include relevant articles from these journals, we exported all articles to InCites, a tool that maintains the research area tags from Web of Science except for articles in the two multidisciplinary areas, which it sorts into more specific Web of Science research areas such as “neurosciences.” Outside the two multidisciplinary areas, the research areas for each article remain the same between Web of Science and InCites.

We narrowed down the pool of articles to include only articles in the research areas listed above, leaving 1,341 articles remaining. Because of the large number of different Web of Science research areas related to psychology, we grouped the articles in those areas (psychology, applied psychology, biological psychology, clinical psychology, developmental psychology, educational psychology, experimental psychology, mathematical psychology, multidisciplinary psychology, psychoanalysis psychology, and social psychology) into one collapsed research area, which we called “psychology” for the present study. The research area “neuroimaging,” which was represented by only six articles in the corpus, was grouped with neurosciences. Thus, the final corpus comprised articles from six research areas (Box 2; Table S1): behavioral sciences, clinical neurology, neurosciences, psychiatry, psychology, and substance abuse.

Because there was a large disparity between the research areas with the largest and smallest numbers of articles in the corpus, we used an algorithm (Chung et al., 2025) that calculates and considers the weight of each research area to extract a sample of 200 articles (Table S1). This weighted sampling technique ensured that the joint distribution of research areas was balanced compared with the overall corpus while preserving the relative order of research areas from most to least frequent. One hundred sixty articles were tagged with a single research area and 40 were tagged with multiple research areas (e.g., both “neurosciences” and “behavioral sciences”; Table S1). No articles in the sample of 200 were tagged with more than two research areas.

To retrieve PDFs of each article in the sample, we used EndNote v.21. For PDFs not downloaded through EndNote, we searched PubMed or Google Scholar manually and downloaded the PDFs through the Emory University or Indiana University Bloomington libraries. For articles not available through either library, we used the Indiana University Bloomington InterLibrary Loan program.

As a first step in our coding process, we assessed articles to confirm each met inclusion criteria. We excluded four articles for being in a language other than English; 22 articles for not including a claim of a sex- or gender-dependent effect in the title (i.e., an article contained a search term in the title that did not refer to a sex or gender difference in the effect of another variable); five articles for being the wrong type (two reviews, two unpublished manuscripts, and one qualitative study); and one article for including only women. In total, we excluded 32 articles during the first round of coding. To maintain the same distribution of articles across the six research areas (see Table S1), we replaced each excluded article with another article from the same research area. When an excluded article was tagged with more than one research area, we replaced it with one tagged with the same two research areas. To identify replacement articles, we used a random number generator to prioritize articles within each research area (or set of two research areas, as relevant) for selection as replacements. During a second round of exclusions assessing the 32 new replacement articles, we ultimately excluded four articles for not including a claim of a sex- or gender-dependent effect in the title. We repeated the replacement process to identify four more articles. The four articles selected in this second round of substitution were kept during coding, bringing the total number of articles coded to 200.

### Coding of articles

We based our codes on those developed by Garcia-Sifuentes and Maney (2021) to categorize analytical approaches to sex-based data. For each article in our sample, we first noted the search term in the title (e.g., “sex-specific,” “males but not females”) and the claim being made. We also identified the model organism used in each article (i.e., human participants or non-human animals such as rats or mice).

Next, we determined the analytical approach the authors used to support the claim in the title (Table 1). We assessed each analytical approach by determining whether authors statistically compared the effect in the title between the sexes. When the claim in the title was too vague to determine what type of test would support it (e.g., the dependent variable was not clearly articulated), we used the abstract to determine what the claim was. In some cases, the abstracts contained additional claims of sex-dependent effects not mentioned in the title; we did not assess analytical approaches for those claims.

We ascertained analytical approaches using information in the Methods and Results sections of each paper, including supplementary material when relevant. Each article was assigned to at least one of the codes listed in Table 1. We assigned articles containing a mix of appropriate (see Table 1, Code 1.1) and inappropriate approaches (see Table 1, Codes 1.2 – 2.3) to the relevant codes comprising inappropriate approaches, as our goal was to track the inappropriate approaches. That is, only articles that took no inappropriate approaches to support the title claim were coded as appropriate.

To assign codes for each article, we first asked whether the effect of interest was compared statistically between females and males directly — that is, sex was incorporated into the statistical model in a way that allowed statistical comparison of the effect of interest between the sexes. We coded articles meeting this criterion as General Analytical Approach #1 in Table 1. When authors tested the claim in the title statistically, they typically did so by testing for an interaction between sex and the other factor of interest (e.g., treatment) using a multifactorial analysis of variance (ANOVA). We considered this type of ANOVA — whether two-way, three-way, or having more factors — as having tested for interactions with sex so long as sex was one of the factors in the model. Occasionally, authors statistically compared the effect between males and females by employing other tests of interaction or moderation or a statistical comparison of within-sex associations (e.g., statistically comparing two within-sex correlations). We coded all such approaches into General Analytical Approach #1.

After establishing that the effect in question was statistically compared across sex, we then noted whether the authors reported the result of the comparison and if so, whether the result indicated a significant difference (e.g., significant sex by treatment interaction in an ANOVA). We assigned articles to one of five codes. We used the first code (see Table 1, Code 1.1) if the *p* value for the interaction or other relevant statistic was reported and was below alpha (we considered alpha to be 0.05 unless indicated otherwise in the article). We considered only articles in Code 1.1 to contain evidence sufficient to support the claim in the title; for the purposes of graphing the results, this code was designated “appropriate” (see Table 1, Evidence designation for graphs). We sorted the remaining articles coded under General Approach #1 into one or more of the four codes 1.2 – 1.5 (see Table 1): the interaction was reported to be significant, but the *p* value or other relevant statistic was not presented (Code 1.2); the authors claimed that they tested for interactions and/or performed ANOVA with sex as a factor but the results of those tests were missing (Code 1.3); the *p* value for an interaction with sex was reported and was above alpha (Code 1.4); the interaction was reported to be non-significant but the *p* value was not reported (Code 1.5).

In addition to the above codes, we also noted whether authors appropriately conducted post-hoc tests of the treatment effect within each sex. We considered post-hoc tests within each sex appropriate only if authors first tested statistically for interactions between treatment and sex, such as with an ANOVA, and reported a significant interaction (Rich-Edwards & Maney, 2023). Therefore, we noted whether authors appeared to conduct within-sex post-hoc analyses regardless of the outcome of interaction tests. This code was not exclusive of the other codes.

When the effect in the title was not statistically compared between females and males, we assigned the article first to General Approach #2 (see Table 1), then into one of three codes: the analytical approach was limited to within-sex analyses, with no relevant between-sex statistical comparisons that would support the claim in the title (Code 2.1; DISS error; see Maney & Rich-Edwards, 2023); the analytical approach included a test for whether sex explained variation in the overall model (e.g., main effect) before within-sex analyses (Code 2.2); no quantitative comparisons were performed in the study at all, despite claims of a sex-dependent effect in the title (Code 2.3).

In some cases, in addition to the DISS error, authors compared the sexes within levels of the other factor — for example, they looked for sex differences within a treatment group while ignoring the control group, without first conducting tests for interactions. Such errors were noted during the coding process.

Two coders (MTO and DLM) independently assessed each article. These coders chose the same codes, without discussion, ∼ 70% of the time. In the cases of discrepant initial codes or uncertainty on the part of either coder, the coders discussed the article until reaching agreement. For eight articles, the best code for the analytical approach remained unclear after our discussion; in these cases, AWB read the article and provided input.

### Journal and article metrics

To assess the relationship between the appropriateness of analytical approaches and the potential impact of the article, we took note of several metrics. First, we used Journal Citation Reports (Clarivate Analytics), which provides the journal’s quartile ranking (Q1, Q2, Q3, or Q4) based on Journal Impact Factor. Because journals are given a new quartile ranking each year, we used the quartile the journal held the year the sampled article was published. Some journals did not have a quartile ranking in their citation reports for the year we were interested in; in this case, we excluded the sampled article from our analysis of the quartiles.

Journal Impact Factors and the associated quartiles are calculated within a journal’s Web of Science subject category and indices, including the Science Citation Index Expanded (SCIE) and the Social Sciences Citation Index (SSCI). A journal tagged with a single research area could have up to two quartile measures, for example, one for SSCI and the other for SCIE. Likewise, a journal tagged with two research areas could have up to four quartile measures. When journals were tagged with additional, non-focal research areas that were not relevant to our study (e.g., developmental biology or endocrinology & metabolism), the quartiles associated with these research areas were not considered in our analysis. Twenty-one articles had discrepant quartiles either because they were tagged with multiple focal research areas (see Table S1, Supplementary Data File 1) or because the quartile in one or more relevant areas was calculated in different indices, SCIE or SSCI. To make the graphs in Fig. 3a and S2a as relevant as possible to neuroscientists, we used the quartile for neuroscience when applicable (n = 15), SCIE over SSCI (n = 5), and clinical psychology over developmental psychology (n = 1). In our analysis of the data (below), we accounted for all quartiles assigned to a given article.

Second, we took note of each article’s Category Normalized Citation Impact (CNCI), obtained from InCites. CNCI is an article-level citation metric that represents the ratio of an article’s observed citation count to the expected citation count for publications of the same document type, publication year, and Web of Science subject category (Clarivate, 2025). We obtained article-level CNCI values by exporting DOI-identified records from Web of Science Core Collection to InCites, then downloading document-level metrics. We used the CNCI value assigned by InCites to each sampled article on Jan. 7, 2026 (InCites dataset updated Dec. 10, 2025; included Web of Science content indexed through Nov. 11, 2025).

### Shared authorship among papers in the sample

We inspected the author lists of all 200 articles to identify potential overlap. Authors with identical full names were confirmed to be the same person by looking at affiliations. Out of 200 articles, 30 shared at least one author with at least one other article in the sample. Eight of these 30 articles appeared in two different pairs and one appeared in three pairs, leading to a total of 20 unique pairs. Based on the sample distribution of the four analytical approaches (appropriate, missing, contradictory, and no direct test) over the entire sample of 200 papers, the probability that a randomly selected pair of articles shared an approach was 40.2%. In the sample with shared authors, eight of 20 pairs (40.0%) shared an analytical approach, a proportion similar to that in the overall sample of articles. Thus, we have no evidence that sharing an author increased the chances of also sharing an analytical approach, and we therefore felt comfortable treating articles as independent observations.

### Data analysis

To test whether the use of appropriate statistical evidence is more common in any area compared with research outside that area, we fit a multivariable logistic regression model including all six research areas with an intercept-only logistic regression model. Because research areas were not mutually exclusive and the same article could be tagged with two, the categorical variable was represented using six binary indicator variables corresponding to the six research areas. In the full model, the odds ratio for each research area represents the change in the odds of an article containing only appropriate statistical evidence when it is tagged with that research area compared with articles not tagged with that research area. The same procedure was followed to test whether the use of the DISS approach was associated with a given research area.

To evaluate the association between model organism (human vs. non-human) and the presence of appropriate statistical evidence to support the claims in the title, we fit a univariate logistic regression that included a single indicator for non-human studies. After excluding four articles that were labeled as both human and non-human, the categories of ‘non-human’ and ‘human’ were mutually exclusive. The estimated odds ratio in this model represents the relative odds of an article on non-humans containing appropriate statistical evidence compared with articles on humans. The same procedure was followed to evaluate the association between model organism and the DISS approach.

To examine associations with quartile, we fit a cumulative link mixed-effects model with each article’s quartile, which articles inherit from the publishing journal, as an ordinal response and the presence of appropriate statistical evidence or DISS error as primary predictors in separate models. Because some of the journals were assigned multiple quartiles from different research areas and indices (SSCI or SCIE), a random intercept for each article was included to account for the within-article correlation. Index was included as a fixed effect. To evaluate the association between the analytical approach and the article’s impact, we fit a linear regression with each article’s CNCI as the response and the presence of appropriate statistical evidence or DISS error as the primary predictor. Log transformation was performed for CNCI prior to the model fitting to improve the normality of the residuals given the right-skewness and strict-positive nature of this measurement. Model coefficients were reported in the format of sympercent (Cole, 2000).

### Assessment of claims over time

We examined how the prevalence of title claims of sex-dependent effects has changed over time in the behavioral and brain sciences. We used Web of Science and InCites, as described above, to search for articles published between 2005 and 2025 that were tagged with at least one of our six focal research areas (Box 2) and included one of our search terms (Fig. 1a) in the title. To illustrate how the rate of sex-dependent claims in our focal research areas changed in relation to other disciplines over time, we repeated the above process for articles tagged with 55 research areas that are biomedical in nature but do not focus on the brain or behavior specifically (e.g., oncology, gerontology, hematology; see Table S3). Research areas that did not relate to medicine, such as political science, geology, and ecology, were not included in our analysis. Articles that were tagged with one or more of our focal research areas as well as one or more of the 55 non-focal research areas were included in the relevant focal area(s) but excluded from the total count in the non-focal area(s) to prevent articles from appearing in both the focal and non-focal counts in the graph. This process applied as well to articles tagged with “multidisciplinary sciences” and “medicine general internal,” both of which were among the non-focal research areas. All articles with these two multidisciplinary tags were exported to InCites and moved to the relevant focal area if tagged by InCites as belonging to a focal area. All searches were conducted on January 10, 2026.

## Acknowledgments

The authors are grateful to Katherine Mao and Kim Powell for help with data collection, to Megan Massa for feedback on an earlier version of the manuscript, and to Yuk Fai Cheong for helpful conversations about data analysis. Figure 2a was prepared with assistance from Chris Goode.

## Author contributions

DLM and MTO conceived, designed, and directed the project. MTO generated the corpus with assistance from CJV, and SC wrote the algorithm used to generate the final sample. DLM and MTO coded the articles with input from AWB. SC analyzed the data under the supervision of AWB. DLM and MTO wrote the manuscript and prepared the tables and figures. All authors assisted with editing the manuscript, tables, and figures and approved the final manuscript. Funding obtained by DLM, AWB and CJV.

## Data availability

All data supporting the findings of this study can be found in Supplementary Data Files 1 and 2. Those files, plus the code used for data analysis, are available at https://osf.io/59djk.

## Footnotes

1 We have not provided citations for quotes because our intention is to provide examples of misunderstandings and unsupported conclusions, not to credit particular authors with such.

