## Supplementary Material for "The “sex-specific effect:” Evaluating analytical approaches to sex-dependence in the behavioral and brain sciences"

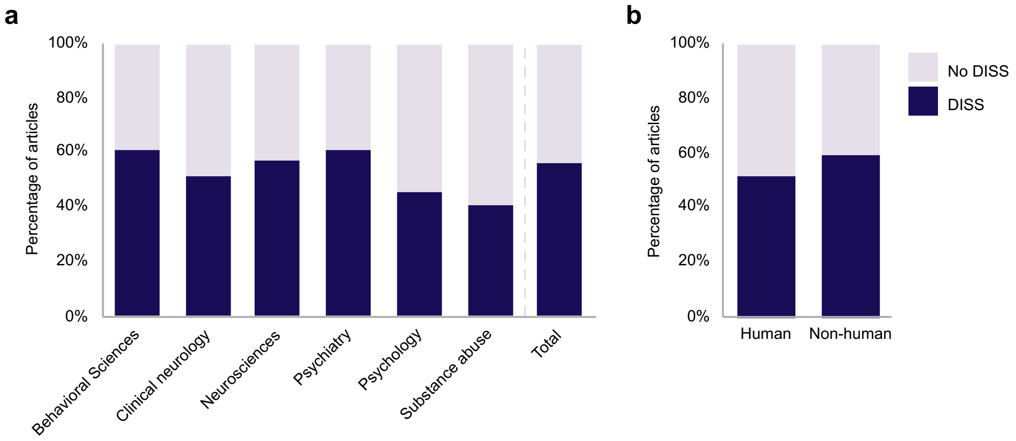


**Fig. S1. Prevalence of DISS approaches across research areas and model organisms.** (a) Across the InCites research areas, articles in the field of psychology and substance abuse were the least likely to feature DISS-based evidence of the claim of sex-dependent effects in the title. Analytic approaches in the “no DISS” category included appropriate as well as inappropriate, non-DISS approaches (see Fig. 2b). Psychology articles were significantly less likely to support the title claim using a DISS approach than non-psychology articles ($p = 0.030$); the same was the case for the area of substance abuse ($p = 0.026$). In the graph, articles that were tagged with two research areas are included in both relevant columns. (b) The prevalence of the DISS error did not differ between studies with only human participants and those with only non-human animals ($p = 0.263$).

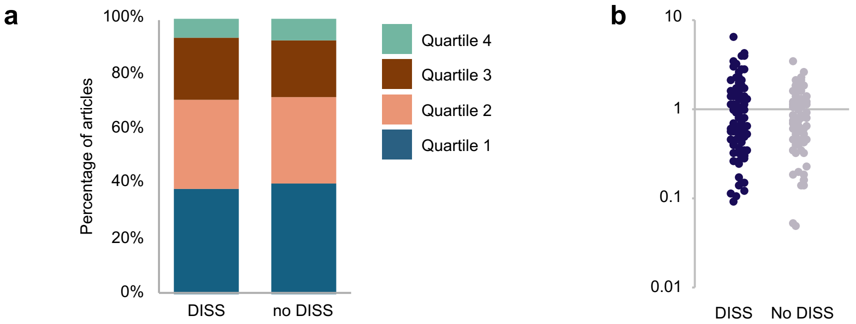


**Fig. S2. Associations between the DISS approach and journal/article impact.** We found no associations between the DISS approach and either (a) the journal’s quartile ranking ($p = 0.590$) or (b) the article’s Category Normalized Citation Impact (CNCI) score ($p = 0.816$). See the caption of Fig. 3 for explanations of these variables.

**Table S1. Distribution of articles in the corpus across InCites research areas.** Numbers in parentheses indicate the number of articles included in the final sample.

| *Research areas* | *2019* | *2020* | *2021* | *2022* | *2023* | *TOTAL* |
|---|---|---|---|---|---|---|
| *Behavioral sciences* | 12 (1) | 9 (1) | 14 (2) | 12 (4) | 14 (2) | 61 (10) |
| *Behavioral sciences; Clinical neurology; Psychiatry* | 0 | 0 | 1 (0) | 0 | 0 | 1 (0) |
| *Behavioral sciences; Neurosciences* | 18 (3) | 19 (1) | 28 (3) | 20 (4) | 20 (1) | 105 (12) |
| *Behavioral sciences; Neurosciences; Psychology* | 1 (0) | 1 (0) | 3 (0) | 1 (0) | 0 | 6 (0) |
| *Behavioral sciences; Psychology* | 7 (1) | 3 (1) | 7 (2) | 5 (1) | 6 (1) | 28 (6) |
| *Clinical neurology* | 17 (6) | 15 (2) | 13 (3) | 13 (5) | 26 (11) | 84 (27) |
| *Clinical neurology; Neurosciences* | 9 (1) | 20 (1) | 15 (0) | 17 (1) | 23 (1) | 84 (4) |
| *Clinical neurology; Neurosciences; Psychiatry* | 1 (0) | 4 (0) | 5 (0) | 3 (0) | 1 (0) | 14 (0) |
| *Clinical neurology; Psychiatry* | 4 (0) | 2 (0) | 5 (0) | 4 (1) | 6 (1) | 21 (2) |
| *Clinical neurology; Neurosciences; Psychiatry; Psychology* | 0 | 0 | 0 | 0 | 1 (0) | 1 (0) |
| *Neurosciences* | 72 (7) | 90 (8) | 97 (11) | 121 (8) | 108 (12) | 488 (46) |
| *Neurosciences; Psychiatry* | 24 (2) | 24 (2) | 26 (2) | 27 (3) | 32 (6) | 133 (15) |
| *Neurosciences; Psychology* | 2 (0) | 2 (0) | 5 (0) | 1 (0) | 4 (0) | 14 (0) |
| *Psychiatry* | 19 (4) | 12 (3) | 20 (8) | 28 (10) | 19 (6) | 98 (31) |
| *Psychiatry; Psychology* | 4 (0) | 6 (0) | 5 (0) | 2 (0) | 12 (0) | 29 (0) |
| *Psychiatry; Psychology; Substance abuse* | 1 (0) | 2 (0) | 1 (0) | 2 (0) | 0 | 6 (0) |
| *Psychiatry; Substance abuse* | 2 (0) | 0 | 4 (1) | 5 (0) | 0 | 11 (1) |
| *Psychology* | 24 (4) | 22 (8) | 20 (3) | 14 (8) | 23 (2) | 103 (25) |
| *Psychology; Substance abuse* | 3 (0) | 3 (0) | 0 | 0 | 1 (0) | 7 (0) |
| *Substance abuse* | 6 (3) | 8 (5) | 15 (4) | 10 (4) | 8 (5) | 47 (21) |
| ***TOTAL*** | 226 (32) | 242 (32) | 284 (39) | 285 (49) | 304 (48) | 1,341 (200) |

**Table S2. Estimated probabilities that an article within a research area would have appropriate evidence.** Probabilities were obtained via marginal predictions from the multivariable logistic regression model.

| *Research Area* | *Probability of Appropriate (95% CI)* |
|---|---|
| *Behavioral Sciences* | 0.199 [0.088, 0.391] |
| *Clinical Neurology* | 0.302 [0.170, 0.477] |
| *Neurosciences* | 0.177 [0.106, 0.280] |
| *Psychiatry* | 0.221 [0.126, 0.359] |
| *Psychology* | 0.387 [0.234, 0.565] |
| *Substance Abuse* | 0.272 [0.127, 0.489] |

Abbreviations: CI = Confidence Interval

**Table S3. Summary of the multivariable logistic regression model assessing the association between six research areas and the use of appropriate evidence**. Each research area was coded as a binary indicator.

| *Research Area* | *OR* | *95% CI* | *p-value* |
|---|---|---|---|
| *Behavioral Sciences* | 1.215 | 0.373, 3.651 | 0.734 |
| *Clinical Neurology* | 3.109 | 0.860, 11.188 | 0.080 |
| *Neurosciences* | 1.346 | 0.467, 3.863 | 0.587 |
| *Psychiatry* | 1.876 | 0.616, 5.569 | 0.258 |
| *Psychology* | 4.705 | 1.255, 18.127 | 0.022 |
| *Substance Abuse* | 2.811 | 0.619, 12.394 | 0.173 |

Abbreviations: CI = Confidence Interval, OR = Odds Ratio

**Table S4. Estimated probabilities that an article within a research area would employ a DISS approach.** Probabilities were obtained via marginal predictions from the multivariable logistic regression model.

| *Research Area* | *Probability of Appropriate (95% CI)* |
|---|---|
| *Behavioral Sciences* | 0.613 [0.420, 0.776] |
| *Clinical Neurology* | 0.515 [0.348, 0.678] |
| *Neurosciences* | 0.572 [0.459, 0.678] |
| *Psychiatry* | 0.615 [0.471, 0.742] |
| *Psychology* | 0.452 [0.289, 0.626] |
| *Substance Abuse* | 0.409 [0.228, 0.618] |

Abbreviations: CI = Confidence Interval, DISS = Difference in Sex-specific Significance

**Table S5. Summary of the multivariable logistic regression model assessing the association between six research areas and employing a DISS approach**. Each research area was coded as a binary indicator.

| *Research Area* | *OR* | *95% CI* | *p-value* |
|---|---|---|---|
| *Behavioral Sciences* | 0.916 | 0.355, 2.388 | 0.855 |
| *Clinical Neurology* | 0.382 | 0.121, 1.162 | 0.093 |
| *Neurosciences* | 0.495 | 0.196, 1.210 | 0.127 |
| *Psychiatry* | 0.687 | 0.268, 1.740 | 0.429 |
| *Psychology* | 0.271 | 0.082, 0.869 | 0.030 |
| *Substance Abuse* | 0.227 | 0.060, 0.824 | 0.026 |

Abbreviations: CI = Confidence Interval, DISS = Difference in Sex-specific Significance, OR = Odds Ratio,

**Table S6. Estimated probabilities of appropriate evidence for human and non-human animal studies, respectively.** Probabilities were obtained via marginal predictions from the univariable logistic regression model.

| *Model Organism* | *Probability of Appropriate (95% CI)* |
|---|---|
| *Human* | 0.344 [0.256, 0.444] |
| *Non-human* | 0.150 [0.092, 0.234] |

Abbreviations: CI = Confidence Interval

**Table S7. Summary of the univariable logistic regression model assessing the association between model organism and the use of appropriate evidence.** Human studies were used as the reference group.

| *Characteristic* | *OR* | *95% CI* | *p-value* |
|---|---|---|---|
| *(Intercept)* | 0.524 | 0.340, 0.792 | 0.003 |
| *Model Organism [Non-human]* | 0.337 | 0.165, 0.663 | 0.002 |

Abbreviations: CI = Confidence Interval, OR = Odds Ratio

**Table S8. Estimated probabilities of a DISS approach in human and non-human animal studies, respectively.** Probabilities were obtained via marginal predictions from the univariable logistic regression model.

| *Model Organism* | *Probability of Appropriate (95% CI)* |
|---|---|
| *Human* | 0.510 [0.411, 0.609] |
| *Non-human* | 0.590 [0.491, 0.682] |

Abbreviations: CI = Confidence Interval, DISS = Difference in Sex-specific Significance

**Table S9. Summary of the univariable logistic regression model assessing the association between model organism and the DISS approach.** Human studies were used as the reference group.

| *Characteristic* | *OR* | *95% CI* | *p-value* |
|---|---|---|---|
| *(Intercept)* | 1.043 | 0.698, 1.559 | 0.838 |
| *Model Organism [Non-human]* | 1.380 | 0.786, 2.435 | 0.263 |

Abbreviations: CI = Confidence Interval, DISS = Difference in Sex-specific Significance, OR = Odds Ratio

**Table S10. Summary of the cumulative link mixed model assessing the association between the use of appropriate evidence and journal quartile rankings.** Journal index indicators were included as covariates for adjustment. The outcome was the journal-specific quartile ranking for each article within each research area, accounting for index, and a random intercept for article ID was specified to account for within-article correlation.

| *Characteristic* | *OR* | *95% CI* | *p-value* |
|---|---|---|---|
| *Statistical Evidence [inappropriate]* | 1.028 | 0.486, 2.177 | 0.942 |
| *Index* | | | |
| *ESCI* | 1.000 | --- | |
| *SCIE* | 0.899 | 0.268, 1.740 | 0.903 |
| *SSCI* | 1.152 | 0.190, 6.973 | 0.877 |

Abbreviations: CI = Confidence Interval, ESCI = Emerging Sources Citation Index, OR = Odds Ratio, SCIE = Science Citation Index Expanded, SSCI = Social Sciences Citation

**Table S11. Summary of the cumulative link mixed model assessing the association between the DISS approach and journal quartile rankings.** Journal index indicators were included as covariates for adjustment. The outcome was the journal-specific quartile ranking for each article within each research area, accounting for index, and a random intercept for article ID was specified to account for within-article correlation.

| *Characteristic* | *OR* | *95% CI* | *p-value* |
|---|---|---|---|
| *Statistical Evidence [inappropriate]* | 0.836 | 0.435, 1.607 | 0.590 |
| *Index* | | | |
| *ESCI* | 1.000 | --- | |
| *SCIE* | 0.890 | 0.159, 4.965 | 0.8894 |
| *SSCI* | 1.120 | 0.184, 6.834 | 0.902 |

Abbreviations: CI = Confidence Interval, DISS = Difference in Sex-specific Significance, ESCI = Emerging Sources Citation Index, OR = Odds Ratio, SCIE = Science Citation Index Expanded, SSCI = Social Sciences Citation

**Table S12. Summary of the linear regression model assessing the association between the use of appropriate evidence and the log-transformed Category Normalized Citation Impact (CNCI).** Entries with CNCI values of zero were omitted because the logarithmic transformation is undefined at zero.

| *Characteristic* | *Sympercent* | *95% CI* | *p-value* |
|---|---|---|---|
| *(Intercept)* | -35.179 | -59.841, -10.518 | 0.005 |
| *Statistical Evidence [inappropriate]* | 2.562 | -25.769, 30.893 | 0.859 |

Abbreviations: CI = Confidence Interval

**Table S13. Summary of the linear regression model assessing the association between the use of appropriate evidence and the log-transformed Category Normalized Citation Impact (CNCI).** Entries with CNCI values of zero were omitted because the logarithmic transformation is undefined at zero.

| *Characteristic* | *Sympercent* | *95% CI* | *p-value* |
|---|---|---|---|
| *(Intercept)* | -31.942 | -48.286, -15.598 | <0.001 |
| *DISS Error* | -2.890 | -25.296, 21.516 | 0.816 |

Abbreviations: CI = Confidence Interval, DISS = Difference in Sex-specific Significance
